# Theoretical prediction of assembly contiguity^*^

**DOI:** 10.64898/2026.11.05.764522

**Authors:** Md Hasin Abrar, Paul Medvedev, Alexei Novikov, Kristoffer Sahlin

## Abstract

Theoretical methods for predicting the performance of assembly algorithms have the potential to guide experimental design on a scale that simulations cannot. A fundamental question of experimental design is how deep to sequence in order to obtain an assembly with long-enough contigs. More precisely, given a certain read length and coverage, what is the expected contig length that an assembler would produce on a given genome? The foundational work of Lander and Waterman tackled this question, but their method is not applicable to modern high-throughput sequencing. In this paper, we derive what we call a Z-value, which is a formula that takes as input the read length, coverage, and basic properties of the genome, and returns a prediction of the mean contig length produced by a simple single-k assembler on error-free data. We show on a benchmark of 14 genomes that the Z-value is highly accurate, with a mean absolute prediction error of less than 2 percent. We further show how this formula can be used to determine the best choice of k for the assembler and how to choose a coverage level that gives contig lengths within a desired fraction of the optimal. Finally, we show how the Z-value can also predict the performance of a more complicated assembler like Velvet and how it can be used to guide the choice between instruments when there is a tradeoff between coverage and read length and the budget is limited.

## 1 Introduction

Sequencing bioinformatics algorithms (i.e. algorithms that analyze sequencing data) are crucial to biological discovery. Their successful development and application rely on the ability to analyze their performance. Such analysis can be done empirically, using real and/or simulated data, or theoretically, using appropriate modeling of both data and algorithms [24]. Empirical analysis is undoubtedly crucial, as theoretical analysis alone cannot capture the complexities of real data. However, unlike theoretical analysis, empirical analysis often fails to provide a deep understanding of performance and can often lead to overfitting [24]. Unfortunately, the toolbox of available theoretical analysis techniques is limited: they are often either too simple to be useful or too complex to be applied. To address this challenge, we seek to develop theoretical techniques that are simple yet predictive of the performance of sequencing bioinformatics algorithms.

Genome assembly with short reads is an excellent testbed for the development of such techniques. It is a mature and stable area with well-understood algorithms and benchmarks. Theoretical analysis has been already applied to develop likelihood-based assemblers [29, 25, 38, 14, 2] and to evaluate assembly quality [39, 35, 11, 16, 13, 22]. There have also been theoretical approaches to evaluate assemblers, either by the conditions under which they can fully reconstruct the original genome sequence [4] or by measuring how much of what is theoretically possible they can reconstruct [37].

In this paper, we target a fundamental question often posed by experimentalists: how much sequencing coverage is needed in order to get an assembly with long-enough contigs? More precisely, given a certain read length and coverage, what is the mean contig length that an assembler produces for a given genome? At what coverage level do the improvements to contig length start diminishing? Unfortunately, our ability to answer these questions is currently limited. Running simulations is often time-prohibitive and, therefore, most labs either take an “overkill” approach, likely wasting resources, or follow the choices made in earlier sequencing projects, likely leading to sub-optimal results. A theoretical predictor of contig length would solve these problems, but previous work is not directly applicable. For example, in a classic paper, Lander and Waterman [21] gave a statistical model on which several follow-up papers improved [33, 40, 41]. However, their approach is only applicable to the pre-high-throughput era, since it is not robust to high coverage and does not account for the fragmentation of contigs due to repeats. Other papers have studied how much coverage and read length is needed to get a single contig [28, 4, 27], but the costs of obtaining a single contig are generally prohibitive and read length cannot be varied in most instruments. Others have characterized assembly contiguity under ideal conditions but do not look at the effect of finite coverage [42, 20].

In this paper, we derive a formula (which we call the *Z*-value) which takes as parameters a read length *L*, coverage level, *k*-mer size, and some limited genome information and gives the predicted mean contig length from a simple single-*k* de Bruijn graph-based assembler. This formula leads directly to a *O*(*L*) time algorithm to compute the mean contig length for a given coverage level, optimizing over all *k*. It also leads to a *O*(*L*) time algorithm to compute the coverage level at which the mean contig length is within a pre-specified factor (e.g. 80%) of the optimal. Using a benchmark of 14 genomes, we show that our predictions are highly accurate, with a mean absolute error of *<* 2%. Furthermore, we show that the *Z*-value can be used to predict the mean contig length of a more sophisticated assembler like Velvet [43]. Finally, we demonstrate how the *Z*-value can be used by an experimentalist with a limited budget to help choose between instruments like MiSeq or HiSeq, where there is a tradeoff between cost and read length.

In Section 2, we give some basic definitions. In Section 3, we describe and motivate the algorithm we will analyze and the model we will use to analyze it. Then, we derive the *Z*-values (Section 4) and the solution to the saturation problem (Section 5). We present our empirical results in Section 6 and conclude with a discussion of limitations and future directions in Section 7.

## 2 Definitions

### De Bruijn graph

A *k-mer* is a string of length *k*. Let *S* be a set of strings; for example, *S* can be a set of *k*-mers, a single chromosome sequence, or the sequences of a multi-chromosomal genome. Given *S* and an integer parameter *k*, the *node-centric de Bruijn graph*, denoted as *dBG*^*k*^(*S*), is a directed graph where the set of nodes corresponds to the set of *k*-mers in *S* and there is an edge between *u* and *v* if the suffix of length *k −* 1 of *u* is a prefix of *v*. A path *v*_1_, …, *v*_*m*_, for *m ≥* 1, in the de Bruijn graph is called a *unitig* if for all 1 *< i ≤ m, v*_*i*_ has one in-neighbor and for all 1 *≤ i < m, v*_*i*_ has one out-neighbor. A unitig is *maximal* if it is not a sub-path of another unitig. The set of maximal unitigs of a de Bruijn graph is unique and is a partition of the *k*-mers of *S* [8]. As we will only be concerned with unitigs which are maximal, we will drop the word maximal and refer to maximal unitigs simply as unitigs. When *S* represents a reference genome assembly, we refer to the unitigs of *dBG*^*k*^(*S*) as *reftigs*. Every reftig is a substring of *S* [23]. Figure 1 illustrates these concepts. For clarity of presentation, we do not consider double-strandedness; however, our experimental results are based on tools [43, 7] that account for double-strandedness by using the bidirected version of de Bruijn graphs [34].

**Figure 1:**
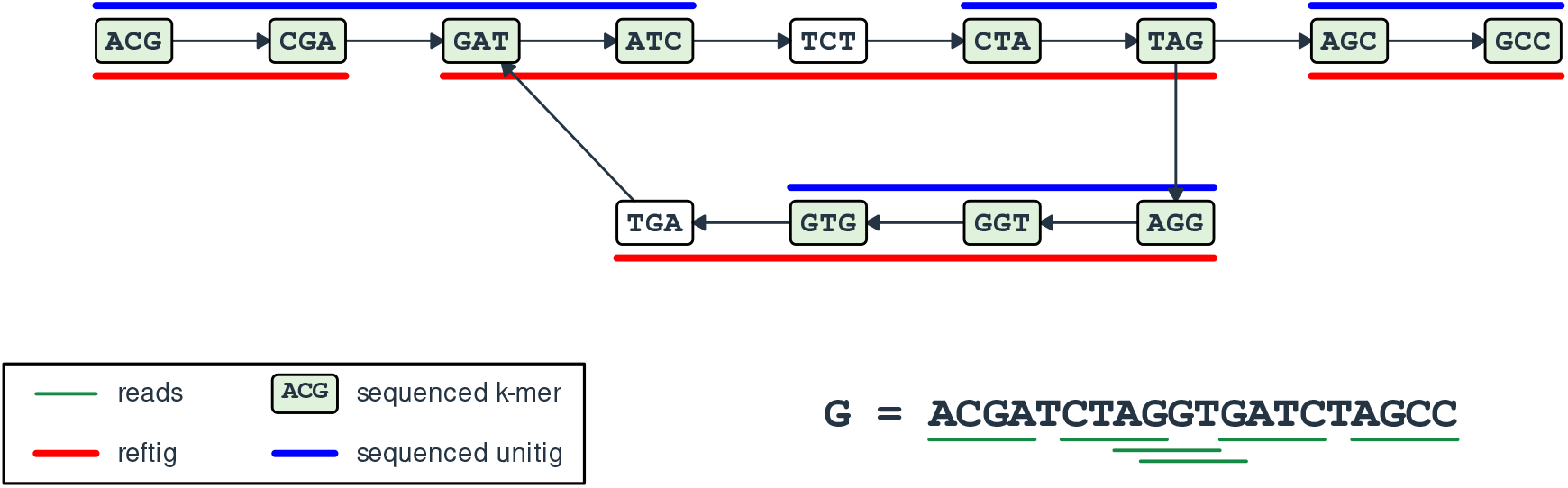
An illustration of de Bruijn graph-related definitions. In this example, there is a single-chromosome genome *G* with a de Bruijn graph for *k* = 3. The reftigs in this graph are shown using red lines (below the vertices). Example sampled reads are underlined in green under *G* and the *k*-mers that are sequenced by these reads are shaded in green. The blue lines (above the vertices) indicate the unitigs in the graph induced by those sequenced *k*-mers, i.e. the de Bruijn graph of the reads. Observe that one unitig is the same as a reftig, two unitigs are substrings of reftigs, and one unitig spans two reftigs (because TGA was not sequenced).

### Sequencing and assembly

Let *G* represent the chromosome sequences of a genome. A *read* is a string generated by taking, uniformly at random, a substring of length *L* from *G*. We define the variable *l* = *L− k*+1 as the *adjusted read length*. A *sequencing experiment* is the result of taking *N* independent reads from *G*. The *depth-of-coverage* (or simply *coverage*) of an experiment is *NL* divided by the number of nucleotides in *G*. At a high level, the *assembly problem* is to take a collection of reads from a sequencing experiment of a latent genome *G* and reconstruct long substrings of *G* [31, 37, 26]. The output of an assembly algorithm is a set of strings *C* called *contigs*.

## 3 Assembly algorithm and analysis model

There are a multitude of algorithms for short-read assembly [32] and choosing which one to analyze is complicated by several factors. The first is that the algorithms differ in their performance, with the Assemblathon2 benchmark [3] finding that there is no clear best assembler. The second is that nearly all the algorithms implement heuristic graph simplification strategies which extend contig length by “collapsing” (i.e. removing) true biological variation. This results in trading-off accuracy for contiguity, but this trade-off relies on ad-hoc heuristics and is therefore difficult to make precise and/or analyze theoretically. Finally, there are other non-standard heuristics that are hard to account for, e.g. error correction and multi-*k* merging.

We therefore consider an idealized assembly algorithm that is designed as a compromise between one whose behavior is amenable to theoretical analysis and one that still captures the basic aspects of state- of-the-art assemblers (pseudocode shown in Alg. 1). This algorithm is based on de Bruijn graphs, as are most state-of-the-art short-read assemblers. For every value of *k*, it constructs the de Bruijn graph from the *k*-mers in the reads. It then measures the mean unitig length of the resulting graph and chooses a value of *k* that maximizes this. The algorithm then outputs the unitigs at this value, which we refer to as contigs since they are the output of an assembly algorithm. We note that our algorithm and its subsequent analysis do not model sequencing error or multi-*k* merging; we leave a detailed discussion of its limitations to Section 7.

### Algorithm 1

A simple assembler

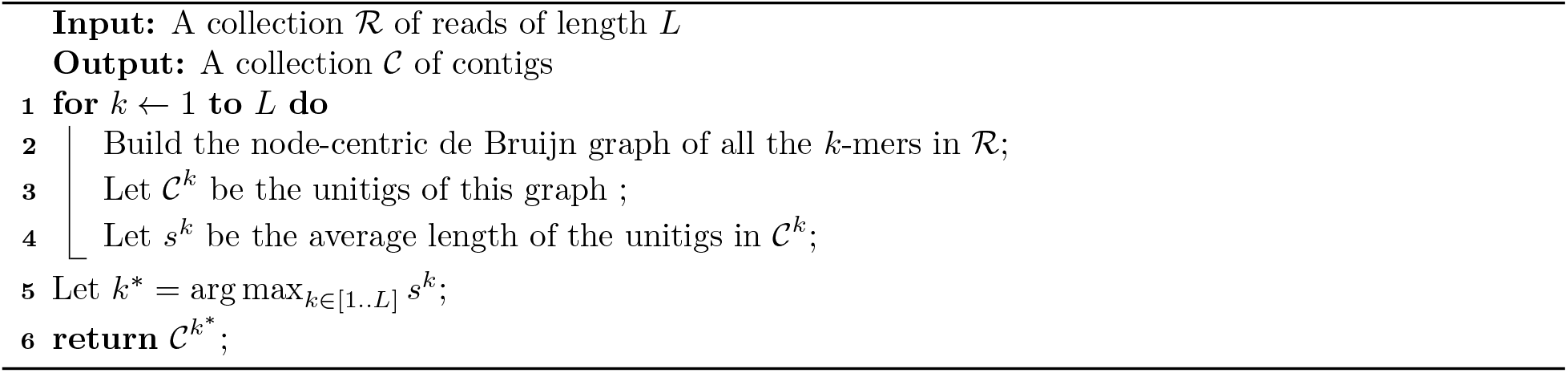

Unfortunately, even for a simple algorithm like Alg. 1, the expected length of contigs remains challenging to analyze. First, for two vertices at a distance *< l* (recall that *l* = *L− k* + 1), their sampling indicators are dependent random variables. These dependencies are not even localized, e.g. when *k < L/*2, every pair of vertices is dependent. Second, a unitig in the *dBG*^*k*^ of the reads may contain an internal vertex that is branching in the *dBG*^*k*^ of the genome (see Figure 1 for an example); as a result, it may not be a substring of the genome. Third, an analysis of Alg. 1 needs *dBG*^1^, …, *dBG*^*L*^; however, this requires prior knowledge of the genome that would be hard to estimate in advance (see Section 7 for a broader discussion of this).

To at least partially side-step these challenges, we adopt a simpler model of a sequencing experiment. Before formally defining it, we present a series of simplifications to provide the intuition. First, we observe that there is a natural relationship between the unitigs *C*^*k*^ of *dBG*^*k*^(ℛ) in Line 3 and the reftigs in *dBG*^*k*^(*G*) (where is the genome being sequenced). Namely, once the coverage of the reads ℛ gets high enough, then *dBG*^*k*^(ℛ) = *dBG*^*k*^(*G*) and *C*^*k*^ are exactly the reftigs in *dBG*^*k*^(*G*). Thus, we can view the reftigs as the limiting bound on the lengths of *C*^*k*^. Building on this, we can think of the underlying genome as having each reftig be its own chromosome, so that sequenced reads are always contained wholly within a reftig. Furthermore, if we drop the first *k*-mer from each reftig-chromosome, both *dBG*^*k*^(*G*) and *dBG*^*k*^(ℛ) will be a disjoint collection of paths. In such a setting, it becomes unnecessary to treat the genome as a string. If the reftig lengths are *r*_1_, …, *r*_*m*_, then we can view the genome as a single interval of length Σ*r*_*i*_, with reftig breaks at positions *r*_1_, *r*_1_ + *r*_2_, *r*_1_ + *r*_2_ + *r*_3_, and so on. Sequencing a read in this setting roughly corresponds to selecting a sub-interval of length *L* from this line; note that this allows read intervals to cross reftig boundaries, but if the reftig lengths are large compared to *L*, then we expect the effect to be negligible. Furthermore, two reads that connect together in the de Bruijn graph are reads whose starting positions are less than *l* bases apart. A unitig in this setting then becomes a maximal interval that does not contain a reftig break and that is bordered on each side by a gap of at least *l* without any read starts.

The second simplification is to allow read locations to be real numbers (rather than integers) and to replace the process of repeatedly drawing *N* locations uniformly at random with a Poisson process that places read starts along the genome at a rate of *N/g*, where *g* is the genome length. We introduce this simplification because it has proven to be useful in studies of contig length in older technologies [21].

Even with these simplifications, we still require prior knowledge of the length of every single reftig in *dBG*^*k*^, for every *k ∈* [1..*L*]. We therefore make a further simplifying assumption: that for a single *k*, the reftig lengths are randomly generated from an exponential distribution with parameter *µ*_*k*_, i.e. with a mean of 1*/µ*_*k*_. Though the reftig lengths observed in practice do not closely follow an exponential distribution, this simplification makes later mathematical analysis more feasible. As a consequence, instead of having reftig breaks as fixed points along the genome line, we can imagine them being generated by a Poisson process with arrival rate *µ*_*k*_. This reduces the *a priori* information we need about the genome to just the average reftig lengths for *k∈* [1..*L*]. After these simplifications, the expected unitig length can be seen as the time from the first read arrival until there is either a reftig break arrival or a stretch of length *l* with no read arrivals. With this intuition in mind, we are now ready to introduce our model formally.

### Model definition

Consider an integer *L >* 1, a sequence of real numbers *µ*_1_ *≥ · · · ≥ µ*_*L*_ *>* 0, and a positive real number *ρ >* 0. We define *read sampling* as a homogeneous Poisson point process with rate *ρ*. We define a family of *L* homogeneous Poisson processes with rates *µ*_1_, …, *µ*_*L*_; each one is referred to as a *k-reftig break* process. We assume that the read sampling process is independent of the *k*-reftig break processes. Let *Y*_first_ be the time of the first read arrival. Let *Y*_last_ be the first time of a read arrival that is followed by an interval of length *l* containing no further read arrivals (note that *Y*_last_ may equal *Y*_first_). For all *k ∈* [1..*L*], let *R*_*k*_ be the first *k*-reftig-break arrival after *Y*_first_. We define the *k-unitig length estimate* as

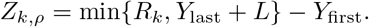

This model is illustrated in Figure 2. As we will see in the following section, this model lends itself to theoretical analysis of contig length while, as we show in the Results, accurately predicting the unitig length in the original de Bruijn graph formulation. Alternative measures of assembly contiguity (e.g. N50) proved more difficult to predict and for Alg. 1, we believe that the mean contig length is a good proxy for the N50.

**Figure 2:**
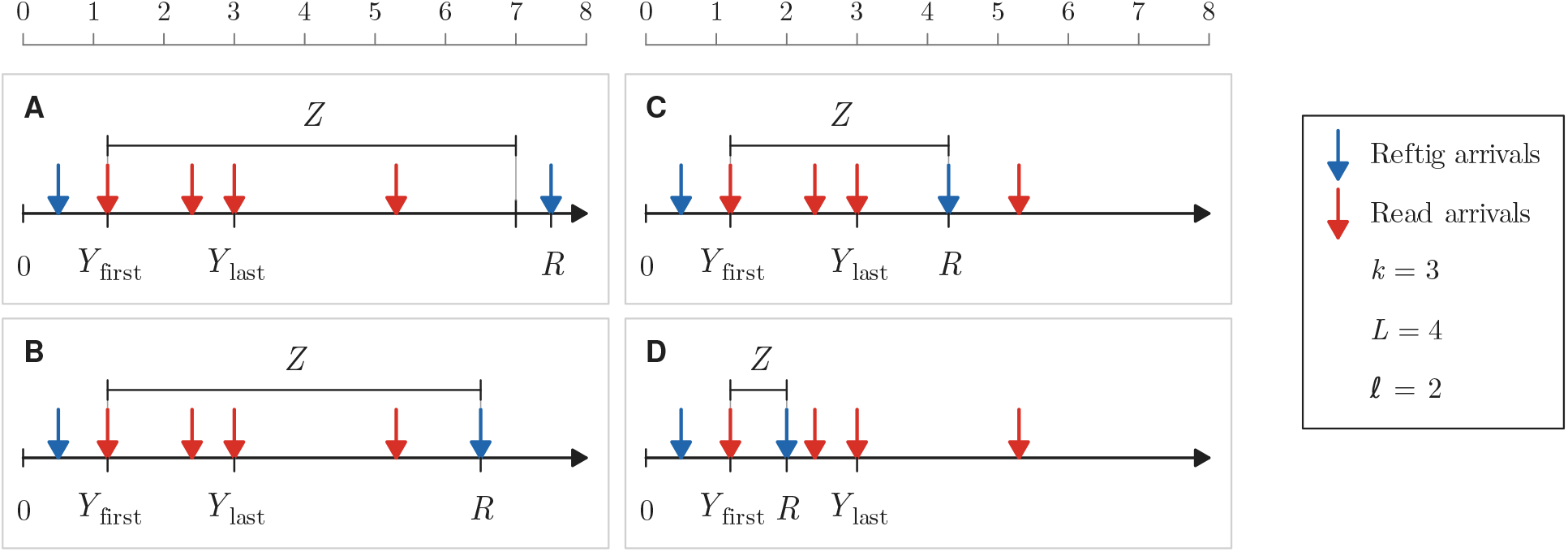
Illustration of the read sampling and *k*-reftig break processes, with *L* = 4 and *k* = 3. *Y*_first_ is the first read arrival, *Y*_last_ is the read arrival preceding the first read-free interval of length *l* = *L − k* + 1, and *R* is the first reftig-break arrival after *Y*_first_. *Z* (shorthand for *Z*_*k,ρ*_) is defined as min*{R, Y*_last_ + *L} − Y*_first_ and reflects the unitig length in this model. The four panels show examples of how various arrival scenarios affect *Z*.

## 4 *Z*-values: mean unitig and contig lengths

In this section, we derive the mean unitig length estimate for a given *k* and *ρ* (i.e. E[*Z*_*k,ρ*_]), the value of *k* estimated to be used by Alg. 1 to achieve the maximum mean contig length (i.e. *k*^*\**^ = arg max_*k∈*[1..*L*]_ E[*Z*_*k,ρ*_]), and the mean estimated unitig length output by Alg. 1 (i.e. 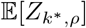). The main technical foundation is the following closed-form formula for E[*Z*_*k,ρ*_].

**Theorem 1**.

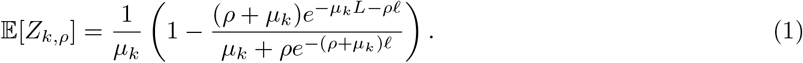

*Proof*. For notational simplicity, define *µ* = *µ*_*k*_ and *Y*_gap_ = *Y*_last_ *− Y*_first_. We can then express *Z*_*k,ρ*_ = min*{R − Y*_first_, *L* + *Y*_gap_*}*. By homogeneity and independence of the reftig break and read sampling Poisson processes, *R − Y*_first_ *∼* Exp(*µ*) and is independent of *Y*_gap_. Recall the property that for an exponentially distributed random variable *T* with parameter *λ* and a positive constant *c*, 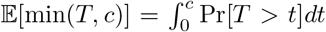. Putting this together and conditioning *Z* on *Y*_gap_ gives

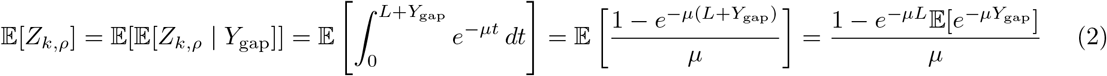

It remains to calculate 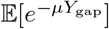. Let *D*_1_, *D*_2_, … be the successive interarrival times after *Y*_first_. These are independent Exp(*ρ*) random variables. Let *N* be the number of read arrivals in (*Y*_first_, *Y*_last_]. Observe that

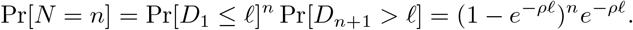

Conditional on *N* = *n, D*_1_, …, *D*_*n*_ remain independent but have an exponential distribution conditioned to be at most *l*. The density of such a distribution is given as *ρe*^*−ρt*^*/* Pr[*D*_*i*_ *≤ l*] on the interval [0, *l*]. Putting this together,

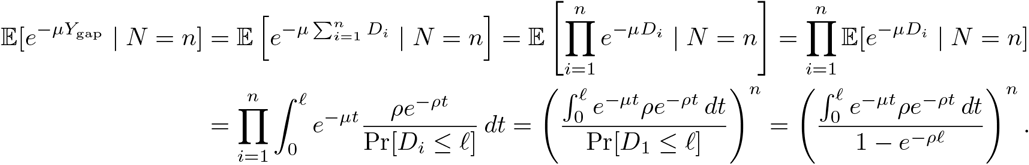

Substituting these expressions into the sum over *n* gives

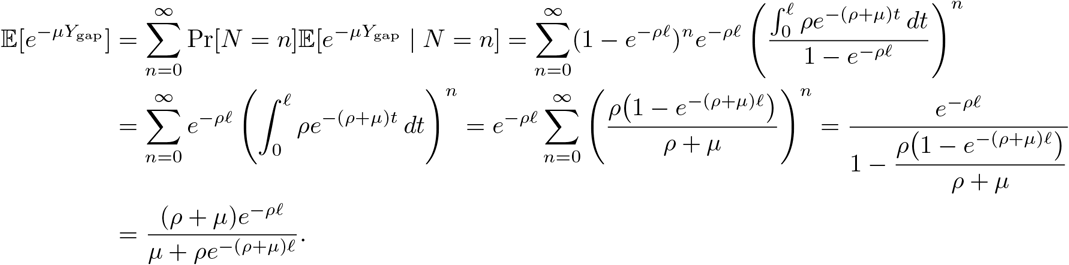

Plugging this into Eq. 2 proves Eq. 1.

We will use *Z*_*k,ρ*_*-value* to refer to the right hand side of Eq. 1, in order to avoid having to always explicitly mention the expectation.

Next, we wish to find the optimal *k* for a given *ρ*, i.e.

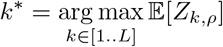

A simple brute force computation over *k∈* [1..*L*] suffices, since a single E[*Z*_*k,ρ*_] evaluation takes constant time using Eq. 1 and doing it for all possible *k* takes only *O*(*L*) time. This computation also gives the estimated mean contig length output by Alg. 1, i.e. 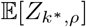. We will use *Z*_*ρ*_*-value* to refer to the result of this computation, i.e. 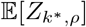.

## 5 Saturation

How much coverage (i.e. how large of *ρ*) is needed to maximize the mean length of the contigs output by Alg. 1? First, observe that there is a natural bound of 1*/µ*_*L*_ on the mean contig length as the coverage goes to infinity. In our simplified model, this bound follows by applying a limit to Eq. 1 and getting lim_*ρ→∞*_ max_*k∈*[1..*L*]_ E[*Z*_*k,ρ*_] = max_*k∈*[1..*L*]_ 1*/µ*_*k*_ = 1*/µ*_*L*_. In the original de Bruijn graph formulation, as coverage goes to *∞*, every *k*-mer, for *k∈* [1..*L*] is sampled, and hence Alg. 1 will choose *k* = *L* and the contig lengths become the *L*-reftig lengths.

Since experimentalists are unlikely to expend large amounts of resources to achieve the maximum contig length, we take the approach that they want to sequence just enough to get to a fraction *α* of this optimal value, e.g. *α* = 80%. This leads to the formal definition of the saturation coverage problem.

### Saturation coverage problem

Given an integer *L >* 1, real numbers *µ*_1_ *≥ · · · ≥ µ*_*L*_ *>* 0, and a real number *α ∈* (0, 1), find

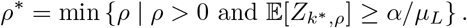

We call *ρ*^*\**^ the *saturating coverage* and say that *k* and *ρ satisfy the saturation criterion* if E[*Z*_*k,ρ*_] *≥ α/µ*_*L*_. Our algorithm for this problem is based on the fact that if we fix *k*, we can derive a formula for 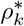, the smallest *ρ* such that *k* and *ρ* satisfy the saturation criterion. The saturating coverage is then obtained by looping over all feasible values of *k* and returning the minimum 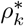, in *O*(*L*) time. The following theorem derives 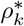 and the feasible values of *k*, and proves the correctness of our algorithm (proof in Appendix).

### Theorem 2

*Let L >* 1 *be an integer, let µ*_1_ *≥ · · · ≥ µ*_*L*_ *>* 0 *be real numbers, and let α ∈* (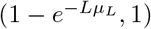) *be a real number. Let k*_*min*_ = min*{k ∈* [1..*L*] *s. t. µ*_*k*_ *< µ*_*L*_*/α}. Then* 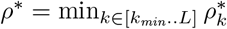, *where*

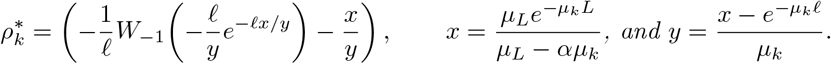

Here, *W*_*−*1_ is the standard notation for the Lambert function [12], which is defined for 1*/e x <* 0 and gives the unique number *y* that is at most −1 and satisfies *ye*^*y*^ = *x*. This function can be evaluated directly in most programming languages.

Note that the condition on *α* in Theorem 2 is not a strong one, e.g. for all the genomes benchmarked in Section 6, *α* can be as low as 5%.

## 6 Empirical results

We have implemented in Python the calculation of *Z*_*k,ρ*_-values, *Z*_*ρ*_-values, *k*^*\**^, and *ρ*^*\**^. ^1^. All runs return results in less than a second and take negligible memory so we therefore do not analyze runtime or memory. The simple assembly algorithm (Alg. 1) as well as the computation of *µ*_*k*_ values is implemented as a trivial wrapper around the bcalm tool [7, 8].

In this section, we demonstrate the accuracy of the *Z*-value and its downstream predictions, compared to actual results of Alg. 1. We also demonstrate how the *Z*-value can be used to predict the performance of the more complex Velvet assembler, and highlight a potential application in choosing a sequencing instrument.

### 6.1 Accuracy

Alg. 1 operates on de Bruijn graphs, but the model we have chosen for analyzing its performance assumes a much simpler framework. We therefore validate the accuracy of the model on 14 reference genomes (Figure 3), computing the *µ*_*k*_ values for *k ∈ {*20, 30, …, 150}. For each coverage *c ∈ {*5, 9, 13, 17, 21, 25}, we generate five simulation replicates of *cg/L* error-free reads of length *L* = 150, where *g* is the genome length (using the ART simulator [15]). We then compute the mean contig lengths output by Alg. 1 for all combinations of *ρ* = *c/L* and *k ∈ {*20, 30, *· · ·*, 150*}*, averaged over the five replicates.

**Figure 3:**
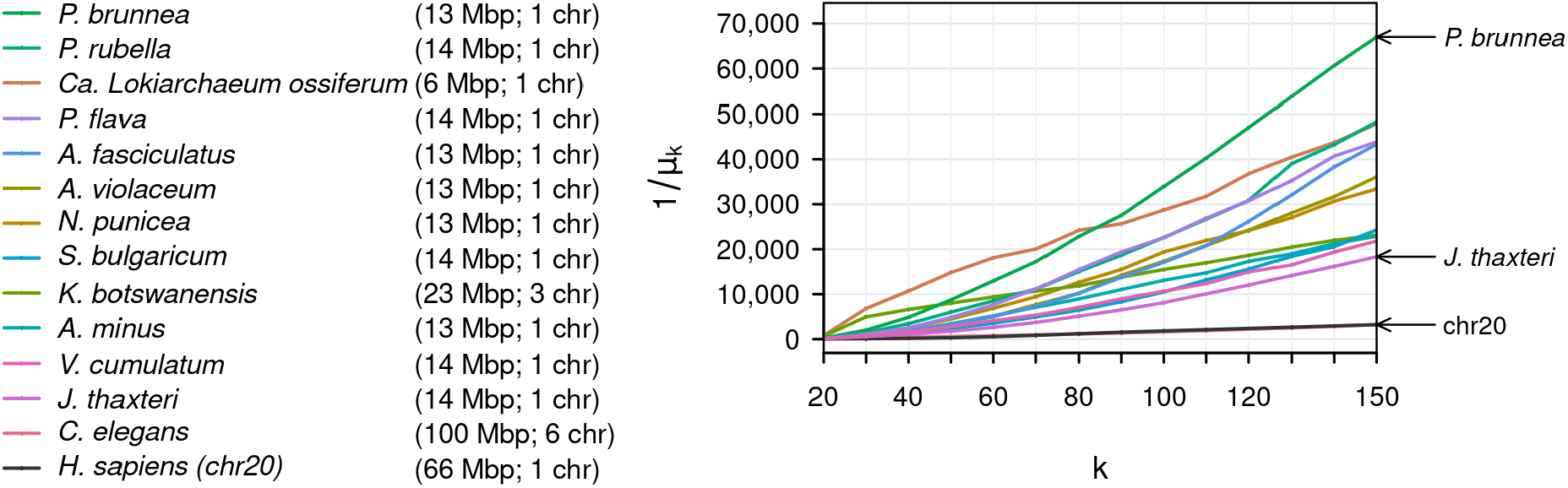
Benchmark genome datasets along with their reftig lengths. All genome assemblies have one contig per chromosome.

Across all 1176 predictions (14 genomes, 6 coverages, and 14 *k* values), the *Z*_*k,ρ*_-value has an absolute prediction error with mean of only 1.7% and maximum of only 8.4%. We also compare the predicted *k*^*\**^ against the optimal choice of *k* by Alg. 1; out of 84 predictions (14 genomes and 6 coverages), 82 were correct, and the other two chose the next best *k*. The *Z*_*ρ*_-value across these 84 predictions has a mean absolute error of only 1.0% against the mean contig lengths returned by Alg. 1, with a maximum of only 3.2%. A full table with these results is included in the appendix (Table S1).

We find this level of accuracy to be remarkable, because Alg. 1 produces contigs that 1) depend on the way that reftigs are connected in the de Bruijn graph; 2) depend on the length distribution of the reftigs, rather than just their mean; 3) can span across multiple reftigs; 4) can be partially covered by reads that start at different reftigs. None of these factors are accounted for in our analysis model, yet the model accurately predicts the unitig and contig lengths.

### 6.2 What can the *Z*-value predict?

To better assess the predictive power of the *Z*-value, we performed deep simulated sequencing of two genomes (*P. brunnea* and *J. thaxteri*), at depths-of-coverage up to 200 *×*. We chose these two because they have different reftig profiles, in spite of having similar lengths (Figure 3).

Figure 4AB show how the *Z*_*k,ρ*_-value behaves as a function of *k* and coverage (while simultaneously re-confirming its accuracy against Alg. 1). These plots confirm patterns already known to assembly practi- tioners, namely that 1) for a given coverage, there is a sweet spot for a value of *k* that maximizes the contig lengths; 2) the sweet-spot-*k* value increases as the coverage increases; 3) the contig lengths increase as the coverage increases. Beyond confirming these patterns, however, the formula gives precise estimates on the unitig lengths as a function of *k* and coverage.

**Figure 4:**
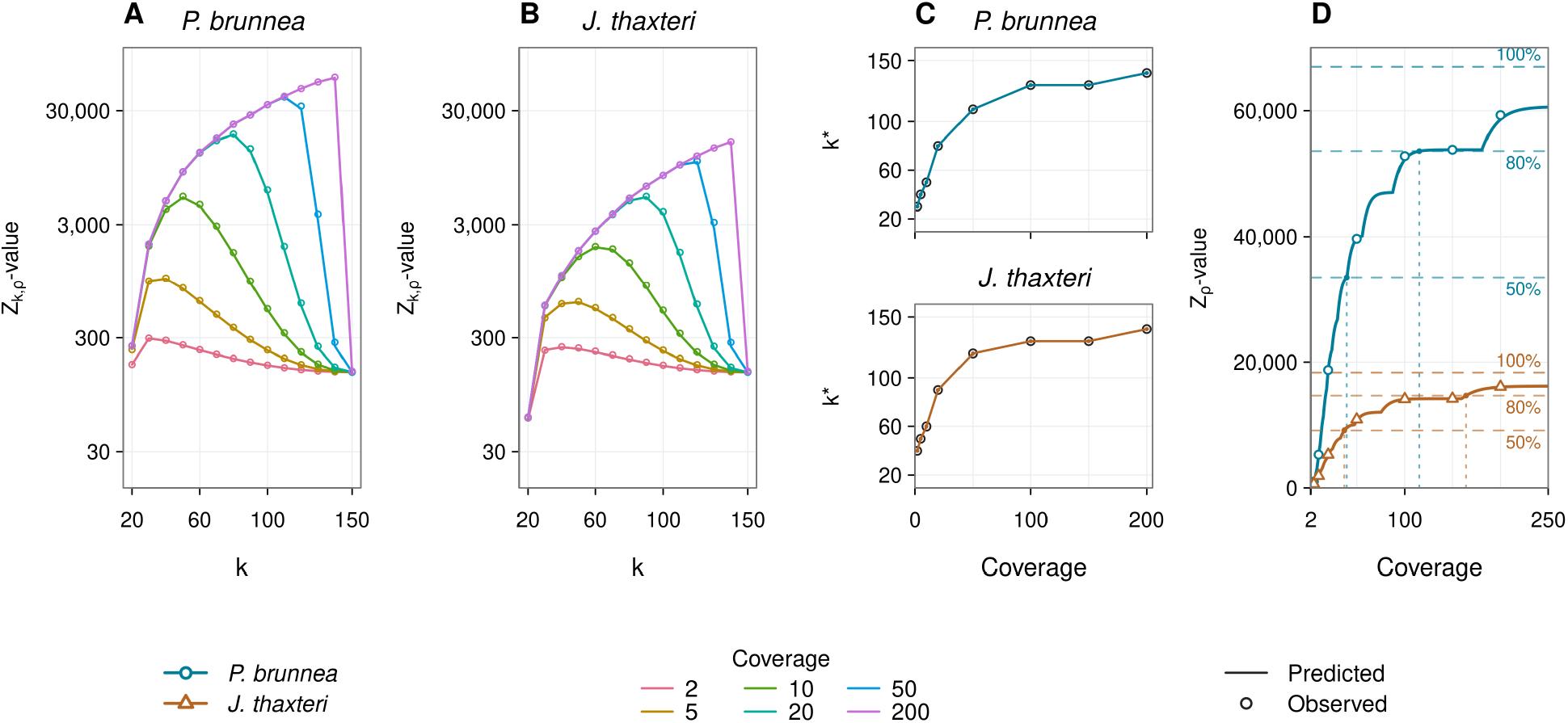
*Z*-value predictions for *P. brunnea* and *J. thaxteri*. Panels (A) and (B) compare *Z*_*k,ρ*_-value predictions (colored lines) with observed mean unitig lengths (open circles). Panel (C) compares the predicted and observed values of *k*^*\**^. Panel (D) shows the *Z*_*ρ*_-value against the mean length of Alg. 1 contigs, with coverage evaluated densely between 2 and 250. Open circles for *P. brunnea* and open triangles for *J. thaxteri* show the Alg. 1 results at the evaluated coverages. The horizontal dashed lines indicate each species’ limiting mean length (i.e. 1*/µ*_150_) and the 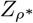-values for saturation coverages *ρ*^*\**^ at *α* = 0.5 and *α* = 0.8.

The *Z*_*k,ρ*_-value gives a simple *O*(*L*) algorithm for computing *k*^*\**^. Figure 4C shows that such predicted *k*^*\**^ values perfectly match the optimal *k* values chosen by Alg. 1. We note that the prediction of the optimal *k* is a side-effect of our model, not its main goal; there are other more advanced methods to select the *k*-mer size for assembly [36, 9].

When sequencing costs are limited, the *Z*_*ρ*_-value can help practitioners predict how limited coverage levels affect the contiguity of the assembly. Figure 4D shows the *Z*_*ρ*_-values as a function of coverage (while confirming agreement with the contig lengths of Alg. 1). For example, if only 100*×* is possible, then the predicted average contig length is 52 Kbp (78% saturation) for *P. brunnea*.

Figure 4D also shows the saturation coverages computed by the algorithm of Theorem 2. For example, for saturation of 80%, the needed coverage is 115*×* (164*×*) for *P. brunnea* (*J. thaxteri*), respectively.

### 6.3 Application: cheap short reads or expensive longer reads?

The *Z*_*ρ*_-value can guide the design of a sequencing experiment beyond just choosing its coverage level. Consider a lab that wants to assemble a genome and must choose between two instruments: *M*_1_, which produces reads of length 300 at a cost of *γ >* 1 monetary units per base, and *M*_2_, which produces shorter but cheaper reads of length 125 at a cost of 1 monetary unit per base. This mirrors a choice between Illumina MiSeq (300 bp reads) and HiSeq 2500 in high-output mode (125 bp reads) [18, 17]. If the lab has a fixed monetary budget, then using *M*_2_ gives *γ* times more coverage than using *M*_1_, but *M*_1_ has longer reads that can span more repeats and thus has longer reftigs. This leads to a trade-off: should the lab choose longer reads or higher coverage?

The *Z*_*ρ*_-value allows the lab to decide on which instrument to choose in order to maximize contiguity, in *O*(*L*) time. Given a budget of *B* monetary units and a genome of length *g*, we can compute the *Z*_*ρ*_-value of *M*_1_ at *ρ* = *B/*300*gγ* and the *Z*_*ρ*_-value of *M*_2_ at *ρ* = *B/*125*g*. Figure 5 shows these plots for different values of *γ∈ {*1, 5, 10}. As expected, for *γ* = 1, it is always preferable to choose the longer reads, regardless of budget. However, when the instruments charge different prices (*γ ∈ {*5, 10}), the plot identifies that there is an intermediate budget range at which higher coverage with shorter reads is better, while outside of this range, lower coverage with longer reads is better. Intuitively, in the lowest coverage range, long reads are better because reads will tend to not overlap and the contigs will be just the reads themselves. At the highest coverage range, the mean contig lengths approach 1*/µ*_*L*_ and hence longer reads are better.

**Figure 5:**
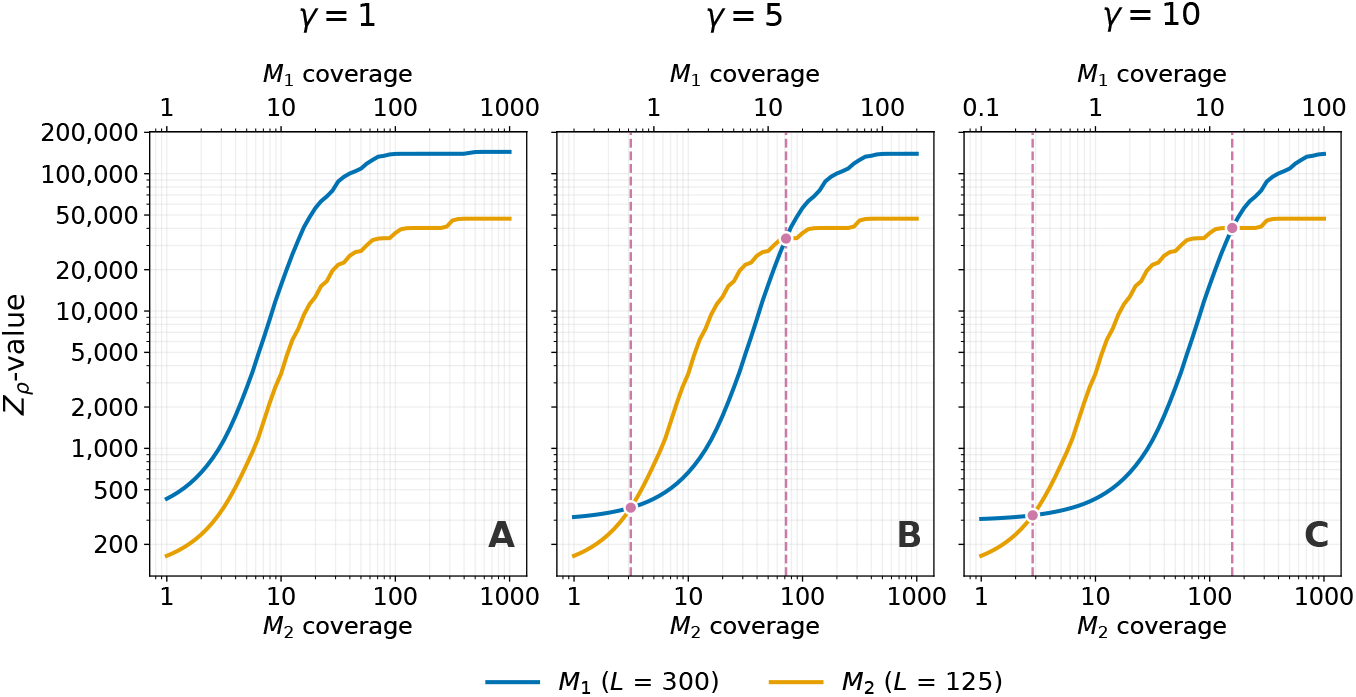
*Z*_*ρ*_-values for *P. brunnea*, as a function of the sequencing budget *B* and choice of sequencing instrument. The blue (darker) curve is an instrument with *L* = 300 at a cost of *γ* monetary units per sequenced base (representing an instrument like MiSeq), and the orange (lighter) curve is an instrument with *L* = 125 at a cost of one monetary unit per base (representing an instrument like HiSeq). The three panels correspond to *γ* = 1, *γ* = 5, and *γ* = 10, respectively. A budget of *B* monetary units buys a coverage of *B/Lγ* on *M*_1_ and *B/L* on *M*_2_. The x-axis is logarithmic in *B* but what is shown at the bottom and top of each panel is the conversion to the respective instrument coverages. Dashed vertical lines mark the budgets at which the choice of instrument changes.

Most relevant for experimental design is the point where this highest coverage regime begins. For example, consider *γ* = 5, which roughly matches the real cost differences between MiSeq and HiSeq. Figure 5B says that if you can afford MiSeq coverage of at least 14 *×*, then you should use MiSeq. Otherwise, if you can afford HiSeq coverage of at least 3 *×*, then you should use HiSeq. Otherwise, you should use MiSeq (though at such low coverages the assembly contiguity is very poor). These numbers are only illustrative, as the trade-offs of these instruments are always in flux.

We note that using simulations to generate Figure 5 would require doing hundreds of assemblies (one per coverage and *k*-mer value at each instrument), each potentially repeated over several replicates. With the *Z*_*ρ*_-value, the figure is generated in *O*(*L*) time, amounting to less than a second.

### 6.4 Extension: predicting the performance of a real assembler

Practical single-*k* assemblers like Velvet [43] include a graph simplification step, i.e. in Line 3 of Alg. 1, they have a more complicated way to compute *C*^*k*^.^2^ More generally, it is challenging to directly model every step that a particular assembler takes. However, these steps do increase assembly contiguity: Figure 6 shows that the mean contig lengths of Velvet are substantially higher than those of Alg. 1. Though this can come with a loss of accuracy, it would nevertheless be interesting to try to predict the performance of such assemblers. To address this issue, we treat Velvet as a black box, and assume that the simplifications that Velvet makes on read data are similar to those it would make on the spectrum of the genome, i.e. on its complete, error-free set of *k*-mers. We therefore introduce a pre-processing step whereby we give the genome spectrum to Velvet and compute the mean contig length of the output, at various *k*-mer sizes. We record the reciprocal of these means as 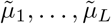 and use them in place of the usual *µ*_*k*_ rates when we do downstream analysis. That is, we compute *Z*-values based on 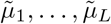 to predict the contig lengths of Velvet on a read set, just as we used *Z*-values based on *µ*_1_, …, *µ*_*L*_ to predict the contig lengths of Alg. 1. In other words, rather than analyzing how Velvet would perform on a reftig distribution of *µ*_1_, …, *µ*_*L*_, we analyze how Alg. 1 performs on a reftig distribution computed with Velvet, i.e. 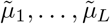.

**Figure 6:**
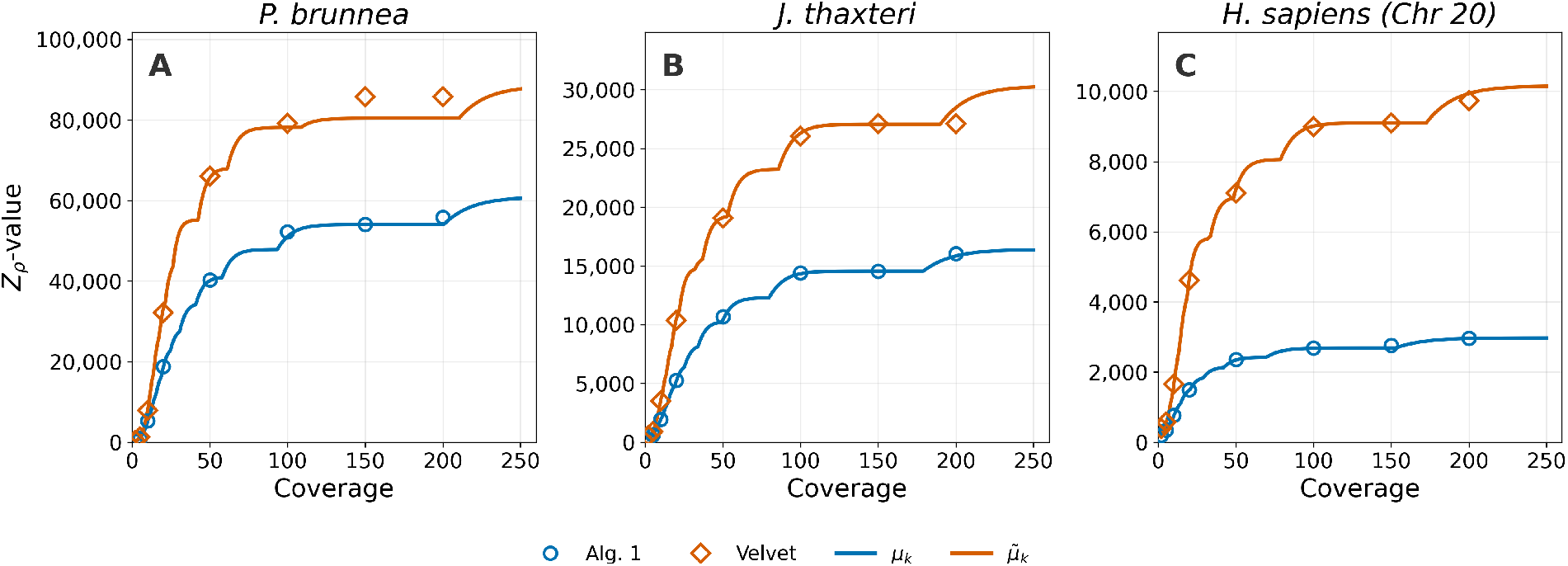
Effectiveness of *Z*_*ρ*_-value predictions for the Velvet assembler, for *P. brunnea* (panel A), *J. thaxteri* (panel B) and human chromosome 20 (panel C). The blue and orange dots show the mean contig length of Alg. 1 and Velvet, respectively, with *L* = 150 and *k ∈ {*21, 31, …, 141*}*. The blue curve shows the *Z*_*ρ*_ values computed using *µ*_*k*_, i.e. the reciprocal reftig lengths from the de Bruijn graph on the full genome. The orange curve shows the *Z*_*ρ*_ values using 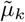, i.e. the reciprocal lengths of contigs output by Velvet when run on the full genome. For each coverage level, five simulation replicates were used and the reported lengths are averages.

Figure 6 shows the performance of this strategy on *P. brunnea, J. thaxteri*, and human chr20. For coverages of 5*×* and above, the *Z*_*ρ*_ predictions using 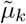 are accurate, with only a 1-3% mean absolute error per species compared to the mean contig lengths computed by Velvet; the maximum error is only 5-6% per species. The predicted *k*^*\**^ is the one chosen by Velvet in all except one case (Velvet chose *k* = 131 instead of *k*^*\**^ = 141 for *J. thaxteri* at coverage 200*×*). These results indicate that our method for accounting for graph simplification allows us to accurately predict the performance of Velvet at coverage of at least 5 *×*.

For the smaller coverage of 2 *×*, the mean error is larger, between 30 and 42% per species. At this coverage, Velvet’s mean contig length is between 252 and 320bp. For such a fragmented assembly and sparse de Bruijn graph, our assumption that the graph simplifications on the full graph mirror that on the sampled graph breaks down. Moreover, Velvet filters out contigs that are shorter than 2*k*, which becomes common at such low coverage. We conclude that our method for graph simplification does not work well when the coverage is too low; however, such low coverage would in any case not be reasonable for performing genome assembly.

## 7 Discussion

In this paper, we derived the *Z*_*k,ρ*_-value, a formula to predict the mean contig length that a simple single-*k* de Bruijn graph assembler achieves, given a certain read length, coverage *ρ* and *k*-mer size (Theorem 1). The formula formed the basis of a simple *O*(*L*) time algorithm that maximizes over all *k* to predict *Z*_*ρ*_, the mean contig length for a given coverage *ρ*, as well as the *k* that achieves it. It also led to an *O*(*L*) time algorithm for computing the saturating coverage, i.e. the smallest coverage needed in order to achieve a mean contig length that is within a given factor of what is possible (Theorem 2). We showed that the *Z*-value and its downstream predictions are highly accurate using a benchmark of 14 genomes, covering various kingdoms, chromosome numbers, genome lengths, and reftig lengths. We also showed that the predictive ability of *Z*-values extends to a real-world assembler that does graph simplifications and other heuristics.

*Z*-values have the potential to do in seconds what it would take CPU-days to do with simulations. To illustrate this, we considered the situation when an experimentalist must choose between using a machine like HiSeq, which has very short reads but cheap per-base cost, and MiSeq, which has longer reads but higher cost. Plotting the *Z*_*ρ*_-values of the two machines as a function of a fixed monetary budget, we could quantify the turning point where it becomes better (with respect to contiguity) to use a MiSeq than a HiSeq.

Perhaps equally important to the predictive algorithms themselves is the model we introduced to do the analysis of the assembly algorithm. The model is a highly simplified view of what happens in a de Bruijn graph assembler, making it possible for us to derive the expected contig size. In spite of these simplifications, the model turns out to be highly accurate. We believe this model may prove useful for predicting the performance of other sequencing bioinformatics algorithms.

Our work has several limitations and natural follow-up directions. The first is the reliance on *a priori* information about the genome to be assembled, namely the values *µ*_1_, …, *µ*_*L*_. Since these are not available in advance of assembling the genome, they should be obtained either through estimation or through the reliance on an already-assembled, similar genome. Estimation may be possible through *k*-mer analysis of a low-coverage sequencing run, e.g. measuring the fraction of *k*-mers that are branching and estimating *µ*_*k*_ based on that (there is some prior work on doing this [36]). Alternatively, if there is a similar genome that has already been assembled, *Z*-values could be computed from the *µ*_*k*_ values of the assembled genome. This is likely to work well if one is assembling a strain of a species with a reference, for example, as is commonly done with *E. coli*. The robustness of this across more distant phylogenetic relationships is a direction of future work.

The second limitation is that our validation with a practical assembler is currently limited to Velvet [43] on error-free reads. For errors, today’s error-rates are very low in short-read instruments and may not have a large effect on the assembly contiguity. On the other hand, assemblers like SPAdes [1] or Minia3 [10, 6] are more widely used today than Velvet. These assemblers use multi-*k* merging, which can sometimes close gaps in a way that a single-*k* assembler cannot. Even though our limitation to single-*k* assemblers limits broader adoption of our prediction algorithms, it represents the first substantial attempt that we are aware of to tackle our prediction question in high-throughput sequencing and forms the basis for future work to increase the breadth of applications.

A natural extension of this work is to long-read instruments such as PacBio or Oxford Nanopore. These instruments have reads that are variable in length (which poses challenges, see e.g. [19]) and have more frequent sequencing errors. Moreover, most popular long-read assemblers (e.g. [5]) are based on string graphs [30] and not de Bruijn graphs. Future work will extend our model to string graphs with reads that are longer, variable in length, and error-prone, allowing us to predict the performance of not only short-read but also long-read assemblers.

## Acknowledgments

PM would like to thank Nithin Varma and Sofya Raskhodnikova for initial work on this problem. This work was supported by NIGMS/NIH grant R01GM146462.

## A Appendix

## Theorem 2

*Let L >* 1 *be an integer, let µ*_1_ *≥ · · · ≥ µ*_*L*_ *>* 0 *be real numbers, and let α ∈* (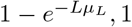) *be a real number. Let k*_*min*_ = min*{k ∈* [1..*L*] *s. t. µ*_*k*_ *< µ*_*L*_*/α}. Then* 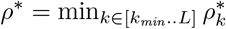, *where*

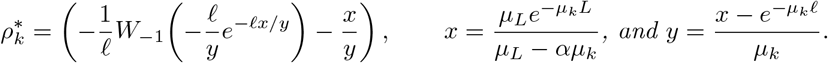

*Proof*. First, observe that for *k < k*_min_, it is impossible to satisfy the saturation criterion. Second, observe that *k*_min_ *≤ L*, hence the set defining *ρ*^*\**^ is non-empty. It remains to show that

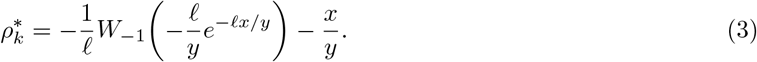

Fix *k∈* [*k*_min_..*L*] and write *µ* = *µ*_*k*_. Define *z* = 1 *−αµ/µ*_*L*_ and observe that the choice of *k* gives *µ < µ*_*L*_*/α* and so *z >* 0. Observe that *x* = *e*^*−µL*^*/z*.

Let *f* (*ρ*) = *e*^*ρl*^ *− x − yρ*. We will show *ρ* belongs to the set defining 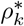 iff *f* (*ρ*) *≥* 0. Starting from Eq. 1,

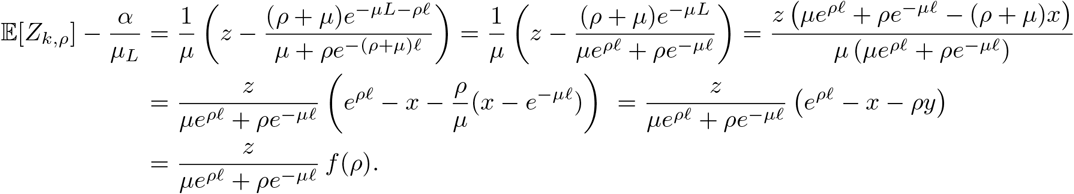

The factor multiplying *f* (*ρ*) is positive, so saturation holds exactly when *f* (*ρ*) *≥*0.

Next, we establish that *f* (0) *<* 0, by showing that *e*^*−µL*^ *> z*, which implies that *x >* 1, *y >* 0, and hence *f* (0) *<* 0. Using the fact that the function (1 *− e*^*−aL*^)*/a* is decreasing for *a >* 0, we get

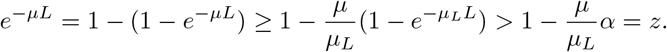

Next, observe that *f* (*ρ*) *→ ∞* as *ρ → ∞* and *f* ^*′′*^(*ρ*) = *l*^2^*e*^*ρl*^ is positive. Thus *f* ^*′*^ is strictly increasing and, since *f* (0) *<* 0, *f* crosses zero exactly once on (0, *∞*) and is nonnegative precisely at and above this crossing. It follows that 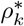 is the unique positive root of *f* (*ρ*).

To find this root, set 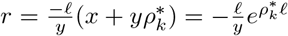. Then,

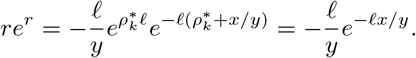

We can solve for *r* explicitly by applying the Lambert function 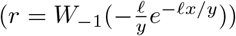 and substituting into 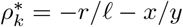 to prove Eq. 3.

To do so, we have to satisfy the conditions of the function. The first is that *r < −*1. This is true because 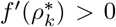 and hence 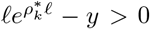 The second is that the argument lies in the domain of *W*_*−*1_, i.e. in [*−*1*/e*, 0).

Set *a* = *lx/y >* 0. Since *x >* 1 and 0 *< ae*^*−a*^ *≤* 1*/e*, we have

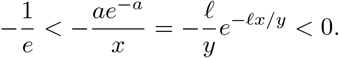

**Table S1:**
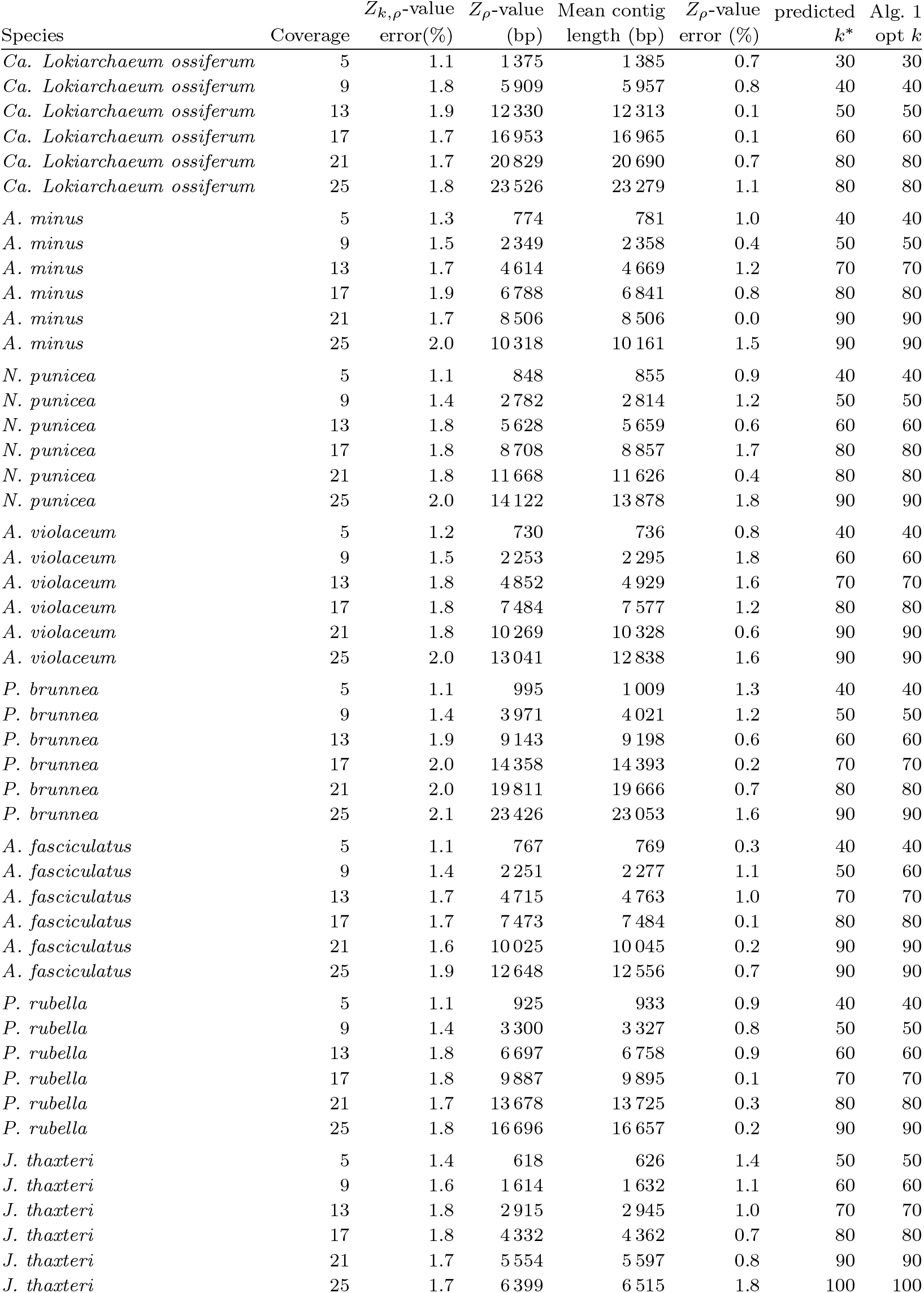
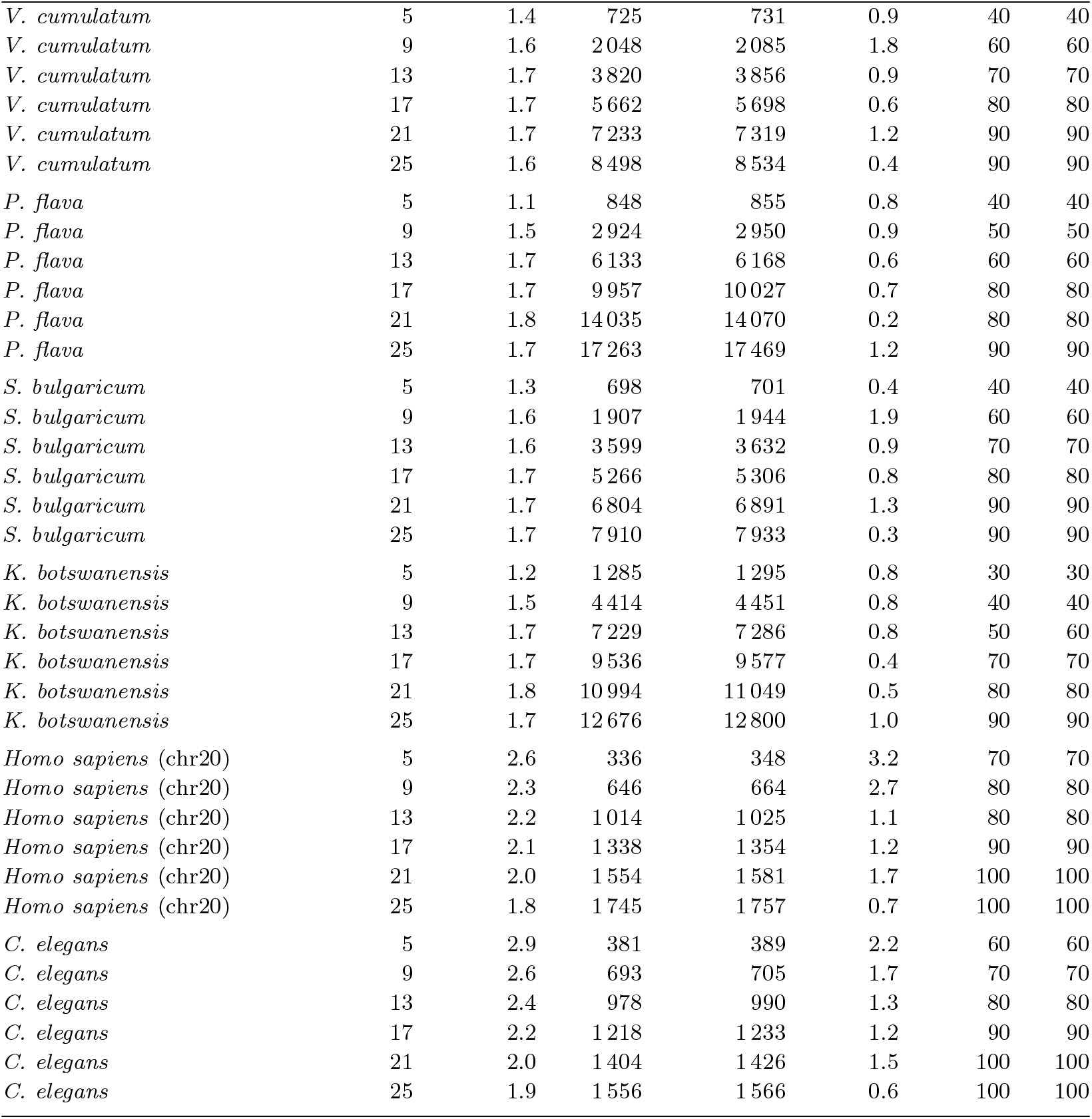
Accuracy of predictions for the 84 genome/coverage settings in Section 6.1. Each row averages the absolute percentage error of the *Z*_*k,ρ*_-value over *k ∈ {*20, 30, …, 150*}*. It also shows the mean contig length of Alg. 1 and the absolute percentage difference between this and the *Z*_*ρ*_-value prediction.

|  |  | *Z*<sub>*k*,*ρ*</sub>-value | *Z*<sub>*ρ*</sub>-value | Mean contig | *Z*<sub>*ρ*</sub>-value | predicted | Alg. 1 |
|---|---|---|---|---|---|---|---|
| Species | Coverage | error(%) | (bp) | length (bp) | error (%) | *k*<sup>\*</sup> | opt *k* |
| *Ca. Lokiarchaeum ossiferum* | 5 | 1.1 | 1 375 | 1 385 | 0.7 | 30 | 30 |
| *Ca. Lokiarchaeum ossiferum* | 9 | 1.8 | 5 909 | 5 957 | 0.8 | 40 | 40 |
| *Ca. Lokiarchaeum ossiferum* | 13 | 1.9 | 12 330 | 12 313 | 0.1 | 50 | 50 |
| *Ca. Lokiarchaeum ossiferum* | 17 | 1.7 | 16 953 | 16 965 | 0.1 | 60 | 60 |
| *Ca. Lokiarchaeum ossiferum* | 21 | 1.7 | 20 829 | 20 690 | 0.7 | 80 | 80 |
| *Ca. Lokiarchaeum ossiferum* | 25 | 1.8 | 23 526 | 23 279 | 1.1 | 80 | 80 |
| *A. minus* | 5 | 1.3 | 774 | 781 | 1.0 | 40 | 40 |
| *A. minus* | 9 | 1.5 | 2 349 | 2 358 | 0.4 | 50 | 50 |
| *A. minus* | 13 | 1.7 | 4 614 | 4 669 | 1.2 | 70 | 70 |
| *A. minus* | 17 | 1.9 | 6 788 | 6 841 | 0.8 | 80 | 80 |
| *A. minus* | 21 | 1.7 | 8 506 | 8 506 | 0.0 | 90 | 90 |
| *A. minus* | 25 | 2.0 | 10 318 | 10 161 | 1.5 | 90 | 90 |
| *N. punicea* | 5 | 1.1 | 848 | 855 | 0.9 | 40 | 40 |
| *N. punicea* | 9 | 1.4 | 2 782 | 2 814 | 1.2 | 50 | 50 |
| *N. punicea* | 13 | 1.8 | 5 628 | 5 659 | 0.6 | 60 | 60 |
| *N. punicea* | 17 | 1.8 | 8 708 | 8 857 | 1.7 | 80 | 80 |
| *N. punicea* | 21 | 1.8 | 11 668 | 11 626 | 0.4 | 80 | 80 |
| *N. punicea* | 25 | 2.0 | 14 122 | 13 878 | 1.8 | 90 | 90 |
| *A. violaceum* | 5 | 1.2 | 730 | 736 | 0.8 | 40 | 40 |
| *A. violaceum* | 9 | 1.5 | 2 253 | 2 295 | 1.8 | 60 | 60 |
| *A. violaceum* | 13 | 1.8 | 4 852 | 4 929 | 1.6 | 70 | 70 |
| *A. violaceum* | 17 | 1.8 | 7 484 | 7 577 | 1.2 | 80 | 80 |
| *A. violaceum* | 21 | 1.8 | 10 269 | 10 328 | 0.6 | 90 | 90 |
| *A. violaceum* | 25 | 2.0 | 13 041 | 12 838 | 1.6 | 90 | 90 |
| *P. brunnea* | 5 | 1.1 | 995 | 1 009 | 1.3 | 40 | 40 |
| *P. brunnea* | 9 | 1.4 | 3 971 | 4 021 | 1.2 | 50 | 50 |
| *P. brunnea* | 13 | 1.9 | 9 143 | 9 198 | 0.6 | 60 | 60 |
| *P. brunnea* | 17 | 2.0 | 14 358 | 14 393 | 0.2 | 70 | 70 |
| *P. brunnea* | 21 | 2.0 | 19 811 | 19 666 | 0.7 | 80 | 80 |
| *P. brunnea* | 25 | 2.1 | 23 426 | 23 053 | 1.6 | 90 | 90 |
| *A. fasciculatus* | 5 | 1.1 | 767 | 769 | 0.3 | 40 | 40 |
| *A. fasciculatus* | 9 | 1.4 | 2 251 | 2 277 | 1.1 | 50 | 60 |
| *A. fasciculatus* | 13 | 1.7 | 4 715 | 4 763 | 1.0 | 70 | 70 |
| *A. fasciculatus* | 17 | 1.7 | 7 473 | 7 484 | 0.1 | 80 | 80 |
| *A. fasciculatus* | 21 | 1.6 | 10 025 | 10 045 | 0.2 | 90 | 90 |
| *A. fasciculatus* | 25 | 1.9 | 12 648 | 12 556 | 0.7 | 90 | 90 |
| *P. rubella* | 5 | 1.1 | 925 | 933 | 0.9 | 40 | 40 |
| *P. rubella* | 9 | 1.4 | 3 300 | 3 327 | 0.8 | 50 | 50 |
| *P. rubella* | 13 | 1.8 | 6 697 | 6 758 | 0.9 | 60 | 60 |
| *P. rubella* | 17 | 1.8 | 9 887 | 9 895 | 0.1 | 70 | 70 |
| *P. rubella* | 21 | 1.7 | 13 678 | 13 725 | 0.3 | 80 | 80 |
| *P. rubella* | 25 | 1.8 | 16 696 | 16 657 | 0.2 | 90 | 90 |
| *J. thaxteri* | 5 | 1.4 | 618 | 626 | 1.4 | 50 | 50 |
| *J. thaxteri* | 9 | 1.6 | 1 614 | 1 632 | 1.1 | 60 | 60 |
| *J. thaxteri* | 13 | 1.8 | 2 915 | 2 945 | 1.0 | 70 | 70 |
| *J. thaxteri* | 17 | 1.8 | 4 332 | 4 362 | 0.7 | 80 | 80 |
| *J. thaxteri* | 21 | 1.7 | 5 554 | 5 597 | 0.8 | 90 | 90 |
| *J. thaxteri* | 25 | 1.7 | 6 399 | 6 515 | 1.8 | 100 | 100 |
| *V. cumulatum* | 5 | 1.4 | 725 | 731 | 0.9 | 40 | 40 |
| *V. cumulatum* | 9 | 1.6 | 2 048 | 2 085 | 1.8 | 60 | 60 |
| *V. cumulatum* | 13 | 1.7 | 3 820 | 3 856 | 0.9 | 70 | 70 |
| *V. cumulatum* | 17 | 1.7 | 5 662 | 5 698 | 0.6 | 80 | 80 |
| *V. cumulatum* | 21 | 1.7 | 7 233 | 7 319 | 1.2 | 90 | 90 |
| *V. cumulatum* | 25 | 1.6 | 8 498 | 8 534 | 0.4 | 90 | 90 |
| *P. flava* | 5 | 1.1 | 848 | 855 | 0.8 | 40 | 40 |
| *P. flava* | 9 | 1.5 | 2 924 | 2 950 | 0.9 | 50 | 50 |
| *P. flava* | 13 | 1.7 | 6 133 | 6 168 | 0.6 | 60 | 60 |
| *P. flava* | 17 | 1.7 | 9 957 | 10 027 | 0.7 | 80 | 80 |
| *P. flava* | 21 | 1.8 | 14 035 | 14 070 | 0.2 | 80 | 80 |
| *P. flava* | 25 | 1.7 | 17 263 | 17 469 | 1.2 | 90 | 90 |
| *S. bulgaricum* | 5 | 1.3 | 698 | 701 | 0.4 | 40 | 40 |
| *S. bulgaricum* | 9 | 1.6 | 1 907 | 1 944 | 1.9 | 60 | 60 |
| *S. bulgaricum* | 13 | 1.6 | 3 599 | 3 632 | 0.9 | 70 | 70 |
| *S. bulgaricum* | 17 | 1.7 | 5 266 | 5 306 | 0.8 | 80 | 80 |
| *S. bulgaricum* | 21 | 1.7 | 6 804 | 6 891 | 1.3 | 90 | 90 |
| *S. bulgaricum* | 25 | 1.7 | 7 910 | 7 933 | 0.3 | 90 | 90 |
| *K. botswanensis* | 5 | 1.2 | 1 285 | 1 295 | 0.8 | 30 | 30 |
| *K. botswanensis* | 9 | 1.5 | 4 414 | 4 451 | 0.8 | 40 | 40 |
| *K. botswanensis* | 13 | 1.7 | 7 229 | 7 286 | 0.8 | 50 | 60 |
| *K. botswanensis* | 17 | 1.7 | 9 536 | 9 577 | 0.4 | 70 | 70 |
| *K. botswanensis* | 21 | 1.8 | 10 994 | 11 049 | 0.5 | 80 | 80 |
| *K. botswanensis* | 25 | 1.7 | 12 676 | 12 800 | 1.0 | 90 | 90 |
| *Homo sapiens* (chr20) | 5 | 2.6 | 336 | 348 | 3.2 | 70 | 70 |
| *Homo sapiens* (chr20) | 9 | 2.3 | 646 | 664 | 2.7 | 80 | 80 |
| *Homo sapiens* (chr20) | 13 | 2.2 | 1 014 | 1 025 | 1.1 | 80 | 80 |
| *Homo sapiens* (chr20) | 17 | 2.1 | 1 338 | 1 354 | 1.2 | 90 | 90 |
| *Homo sapiens* (chr20) | 21 | 2.0 | 1 554 | 1 581 | 1.7 | 100 | 100 |
| *Homo sapiens* (chr20) | 25 | 1.8 | 1 745 | 1 757 | 0.7 | 100 | 100 |
| *C. elegans* | 5 | 2.9 | 381 | 389 | 2.2 | 60 | 60 |
| *C. elegans* | 9 | 2.6 | 693 | 705 | 1.7 | 70 | 70 |
| *C. elegans* | 13 | 2.4 | 978 | 990 | 1.3 | 80 | 80 |
| *C. elegans* | 17 | 2.2 | 1 218 | 1 233 | 1.2 | 90 | 90 |
| *C. elegans* | 21 | 2.0 | 1 404 | 1 426 | 1.5 | 100 | 100 |
| *C. elegans* | 25 | 1.9 | 1 556 | 1 566 | 0.6 | 100 | 100 |

## Footnotes

1 The implementation and reproducibility details are available on GitHub: https://github.com/pashadag/contiguity_predictor

2 Technically, velvet itself does not try multiple *k* values, but for the sake of presentation clarity, we use *Velvet* to refer to the version that runs velvet on all possible *k* values and returns the best result.

